# A novel machine learning based approach for iPS progenitor cell identification

**DOI:** 10.1101/744920

**Authors:** Haishan Zhang, Ximing Shao, Yin Peng, Yanning Teng, Konda Mani Saravanan, Huiling Zhang, Hongchang Li, Yanjie Wei

## Abstract

Identification of induced pluripotent stem (iPS) progenitor cells, the iPS forming cells in early stage of reprogramming, could provide valuable information for studying the origin and underlying mechanism of iPS cells. However, it is very difficult to identify experimentally since there are no biomarkers known for early progenitor cells, and only about 6 days after reprogramming initiation, iPS cells can be experimentally determined via fluorescent probes. What is more, the ratio of progenitor cells during early reprograming period is below 5%, which is too low to capture experimentally in the early stage.

In this paper, we propose a novel computational approach for the identification of iPS progenitor cells based on machine learning and microscopic image analysis. Firstly, we record the reprogramming process using a live cell imaging system after 48 hours of infection with retroviruses expressing Oct4, Sox2 and Klf4, later iPS progenitor cells and normal murine embryonic fibroblasts (MEFs) within 3 to 5 days after infection are labeled by retrospectively tracing the time-lapse microscopic image. We then calculate 11 types of cell morphological and motion features such as area, speed, etc., and select best time windows for modeling and perform feature selection. Finally, a prediction model using XGBoost is built based on the selected six types of features and best time windows. Our model allows several missing values/frames in the sample datasets, thus it is applicable to a wide range of scenarios.

Cross-validation, holdout validation and independent test experiments showed that the minimum precision is above 52%, that is, the ratio of predicted progenitor cells within 3 to 5 days after viral infection is above 52%. The results also confirmed that the morphology and motion pattern of iPS progenitor cells is different from that of normal MEFs, which helps with the machine learning methods for iPS progenitor cell identification.

**Author Summary:** Identification of induced pluripotent stem (iPS) progenitor cells could provide valuable information for studying the origin and underlying mechanism of iPS cells. However, it is very difficult to identify experimentally since there are no biomarkers known for early progenitor cells, and only after about 6 days of induction, iPS cells can be experimentally determined via fluorescent probes. What is more, the percentage of the progenitor cells during the early induction period is below 5%, too low to capture experimentally in early stage. In this work, we proposed an approach for the identification of iPS progenitor cells, the iPS forming cells, based on machine learning and microscopic image analysis. The aim is to help biologists to enrich iPS progenitor cells during the early stage of induction, which allows experimentalists to select iPS progenitor cells with much higher probability, and furthermore to study the biomarkers which trigger the reprogramming process.

## Introduction

Induced pluripotent stem (iPS) cells are cells with embryonic-like state reprogrammed from mouse embryonic or adult fibroblasts by introducing the defined factors[1]. Since Takahashi and Yamanaka[1] first proposed the methods of reprogramming somatic cells to iPS cells, it has become an important method for clinical cell therapy, and revolutionized regenerative medicine[2], such as platelet deficiency[3], spinal cord injury[4], macular degeneration[5], Parkinson’s disease[6] and Alzheimer’s disease[7]. However, obstacles still remain in scientific and clinical applications for iPS cells because of potential tumorigenicity and low efficiency of reprogramming technique[8–10]. Tumorigenicity is attributed to the introduction of tumorigenic factors such as Oct4, Sox2, Klf4 and c-Myc, of which over-expression is generally associated with tumors. Inefficiency concerns low frequency for reprogramming cells, which is less than a small proportion of 5%. In some induction protocols, the ratio of progenitor cells during the early stage of reprogramming is even under 0.5%.

The above-mentioned obstacles are mainly due to poor understanding of molecular mechanisms in iPS cell reprogramming, which ultimately prevented this technology from a wide range of scientific and clinical applications. Theoretical mechanisms models are proposed such as two-step process model[11] and seesaw model[12], most of which focus on how factors such as Oct4, Sox2, Klf4, and c-Myc induce pluripotency. Experimental approaches based on epigenetic profiling, RNA screening or single-cell analysis for uncovering the mechanisms are limited by the low reprogramming efficiency or the lack of biomarkers for progenitor cells [13–20].

Recent studies found that iPS progenitor cells differed from normal MEFs in morphology, motion or proliferation rate. Smith et al.[21] found that iPS progenitor cells showed smaller cellular area and higher proliferative rate than normal MEFs via time-lapse imaging. Zhang et al.[22] also found that iPS cells exhibited distinct morphology features and different proliferative rate comparing with larger and quiescent differentiated cells. Li et al. [23] showed the mesenchymal-to-epithelial transition, a process with significant morphological changes, was a key cellular mechanism for induced pluripotency. Megyola et al.[24] demonstrated that migratory motions for progenitor cells were often distinct in direction and distance to bring distant progenitor cells together. Most of these studies relied on time-lapse microscopy, which allowed studying/tracing cellular events in early reprogramming by direct observation [24]. Since iPS progenitor cells exhibit unique morphology and motion features, computational methods, especially machine learning based methods, could provide an alternative method to identify iPS progenitor cells in the early stage of reprogramming process through learning the morphology and motion patterns of iPS progenitor cells.

Usually cell detection, segmentation and tracking are firstly required for computational methods to study cell images. Li et al.[25] proposed DCELLIQ for cell nuclei tracking based on neighboring graph and integer programming technique. Dzyubachyk et al.[26] relied on coupled active surfaces algorithm for cell segmentation and tracking in time-lapse fluorescence microscopy images. Maška et al.[27] presented a tracking method for fluorescent cells based on coherence-enhancing diffusion filtering and Chan-Vese model. Türetken et al.[28] proposed an integer programming approach for tracking elliptical cell populations in time-lapse image sequences. Payer et al.[29] developed a recurrent fully convolutional network architecture for instance segmentation and tracking with training network using an embedding loss based on cosine similarities.

Recently machine learning/deep learning methods have been extensively developed for the prediction and study of cell images. Using cell images, Erdmann et al.[30] introduced a machine learning based framework for image-based screen analysis. Valen et al.[31] tried to solve cell image segmentation problem utilizing deep convolutional neural networks, and demonstrated its effectiveness in segmenting fluorescent images of cell nuclei. Chen et al.[32] achieved high classification accuracy in label-free white blood T-cells against colon cancer cell via a deep learning method. Similarly with a deep convolutional neural network method, Kraus et al.[33] analyzed the microscopic images for yeast cell and other pheromone-arrested cells, and Gao et al.[34] achieved a high ranking in the human epithelial-2 cell image classification competition hosted by ICPR2014. Together with principal component analysis, machine learning method can be used to infer regulatory network patterns underlying stem cell pluripotency[35]. The ability of machine learning has been demonstrated with its extensive application for cellular image data, however, it has been seldom used in the identification of iPS progenitor cells in the early stage.

In this article, we propose a machine learning based approach to detect iPS progenitor cells during the early stage of reprogramming. Given the cell images recorded via live-cell imaging system during the reprogramming process, the paper aims to identify iPS progenitor cells against normal MEFs in the same stage. Since the iPS progenitor cell to normal MEFs ratio is usually below 5%, this makes the identification problem very difficult. In the paper we use Imaris, a software from Bitplane, to analyze and process microscopic cell images from live-cell imaging system. Surpass, a module of Imaris is then used to extract cell numerical information in the same time period. We then develop a machine learning method for identification of iPS progenitor cells based on the extracted morphological and motional features. The prediction model is built with XGBoost based on the selected six types of features and time windows. In our method, cell division is not considered, and frames contained in selected time windows are uniform. The model performance is evaluated by three different validation methods. When tested on labeled datasets with a ratio of about 1:5 between progenitor cells and normal MEFs, the prediction precision to identify iPS progenitor cells is above 52% during the first 1-3 days of reprogramming after adding iCD1 medium. The image-based machine learning method allows experimentalists to select iPS progenitor cells with much higher probability, and furthermore to study the biomarkers which trigger the reprogramming process.

## Materials and Methods

The workflow used in the paper is presented in **Fig 1**, which mainly includes feature extraction, preprocessing with missing values, feature selection, machine learning for training and validation. In this workflow, we acquire time-lapse images through experiments firstly, then we label iPS progenitor cells and normal MEFs manually to generate datasets by tracing images retrospectively. Next, we generate 11 types of morphology and motion features with Imaris software. After the feature extraction, we perform time window selection and a two-step feature selection. Finally, we build the prediction model based on the selected six types of features and six time windows. The machine learning algorithm for modeling is XGBoost, a gradient boosting tree[36]. In the following sections, we will describe the steps of our model in detail.

**Fig 1.**
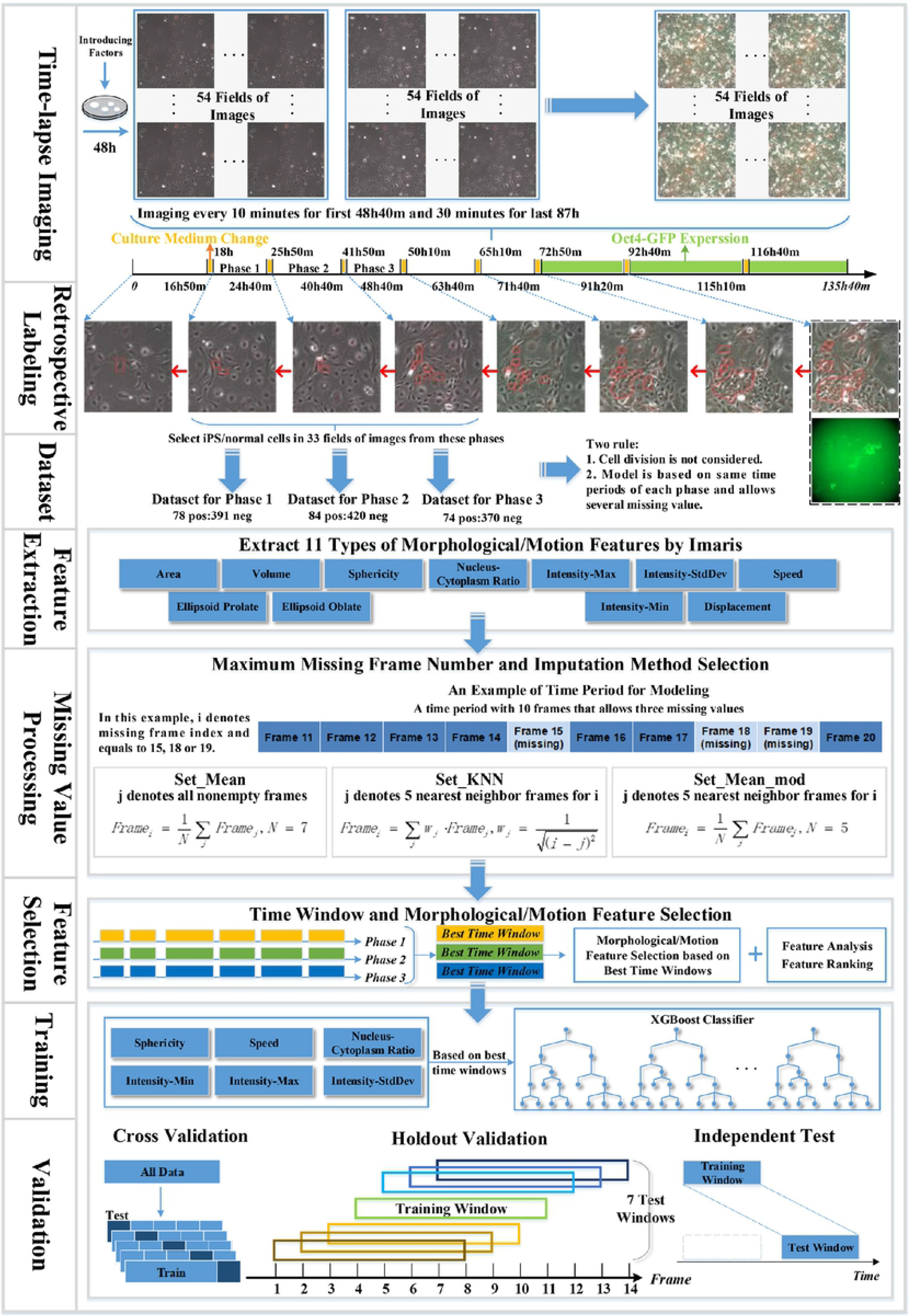
Flow chart of the machine learning based approach for iPS progenitor cell identification. In time-lapse imaging, we record the reprogramming process periodically among 54 fields after 48h of viral infection. For retrospective labeling, the figure only shows the labeled cell images of the first frame of all eight phases. Only datasets from phase 1, 2 and 3 are used for model training and testing.

### Cell culture and generation of iPS cells

Mouse embryonic fibroblasts (MEFs) are derived from E13.5 embryos carrying the Oct4 promoter-driven GFP reporter gene[37] and maintain in DMEM (HyClone) supplemented with 10% FBS (Gibco). To generate iPS cells, MEFs within two passages are seeded at a density of 5×10^4^ cells/well in 6-well plates and cultured overnight. The next day, MEFs are infected with retroviral supernatants containing the DsRed gene and three reprogramming factors (Oct4, Sox2, Klf4) twice in a 48h process. After 48h of infection, iCD1 medium[38] is changed every day to achieve high reprogramming efficiency. iPS cell colonies are obtained 5-7 days post-treatment in iCD1 based on the Oct4-GFP expression.

### Time-lapse imaging

Reprogramming process is recorded using an Olympus IX81 live cell imaging system equipped with a 10× UPlanFL objective, iXon3 EMCCD Camera. The date on which viral supernatants are removed and iCD1 medium are added is defined as Day 0. From Day 0, MEFs images are taken for a total time of 135 hours and 40 minutes. For the first 48 hours and 40 minutes, both bright-field and red fluorescence images are acquired at 10-minute intervals. After two days of the dual-channel imaging, a green fluorescence channel is added to indicate the expression of Oct4-GFP and acquisition intervals are adjusted to 30 minutes. Motorized Stage Control is used to follow cells in the same field and a total of 54 fields are selected at each time for further analysis.

Cell images taken within the first 48 hours and 40 minutes since Day 0 are used to construct the dataset because after this time the Oct4-GFP is added to identify the progenitor cells experimentally and the paper tries to identify/predict progenitor cells using computational methods as early as possible.

### Cell segmentation and numerical feature extraction

The original files are time-lapse microscope images in TIFF format, whose pixels are 770 * 746 and the actual size is 1000 microns * 967 microns. Because some fields do not show distinct Oct4-GFP signals and result in no signals for iPS cells in these fields, we only use images from 33 fields for modeling. Imaris (Version 7) software is used to segment cells in the images of these 33 fields and extract the corresponding numerical features for the segmented cells. During this process, the parameter values of cell and nucleus intensity are set the same for all the cells in each field, and cell tracking duration parameter of greater than 5000s is used. Imaris utilizes red fluorescent channel for cell segmentation and tracking. The image segmentation is based on the Watershed Algorithm, which is very sensitive to weak edges and intensity in images.

Features are computed for each segmented/identified cell image at different time frames by Imaris, and these features denote the morphological and movement information of the segmented cells during reprogramming. Overall 11 types of features are extracted (volume, area, sphericity, ellipsoid-prolate, ellipsoid-oblate, nucleus-cytoplasm volume ratio, displacement, speed, Intensity-stdDev, Intensity-Max, Intensity-Min) and each type contains features in several frames of the selected uniform time windows. The detailed list of features is presented in Part 1 of the S1 File.

### Cell image dataset generation

Cell image datasets for machine learning consist of normal MEFs cell images and progenitor cell images within the first 48 hours and 40 minutes. The datasets will be used by our machine learning method in the training and testing processes.

At first, we manually label iPS progenitor cell and normal MEFs cell images identified by Imaris software within the first 48 hours and 40 minutes. Experimentally iPS cells can be determined only by Oct4-GFP expression signal, which cannot be observed until the seventh day after transfection with Yamanaka’s factors. Cells showing green fluorescence in images are considered as iPS cells. We can then label iPS progenitor cells in the early reprogramming process by cell image backtracking. The corresponding cell images are retrospectively traced frame by frame from GFP expression to the first 48 hours and 40 minutes (**Fig 1**). Due to three one-hour iCD1 medium changes, the total reprogramming period is divided into four periods, the first period is 16 hours and 50 minutes long, from 18 hours to 24 hours and 40 minutes denoted as phase 1 in the paper, the second from 25 hours and 50 minutes to 40 hours and 40 minutes denoted as phase 2, and the third from 41 hours and 50 minutes to 48 hours and 40 minutes denoted as phase 3. In this paper, we focus on these three periods (phases 1, 2 and 3) only because of tiny ratio for iPS progenitor cells in the first 16 hours and 50 minutes, which is even less than 2%.

Two rules are applied in the paper for generating the cell image datasets, (1) cell division is not considered; (2) frames from the same window of each phase are selected for modeling among uniform time periods. When cell division is taken into account, features in the mother cell and its daughter cells are not comparable, for example, the area of mother cell is much bigger than that of its daughter cells, thus the machine learning model will fail to process this cell. The second rule guarantees that time dimension (time period and length) for the cell image data samples should be uniform.

For each cell, not every image in different frames can be identified by Imaris due to the fact that different parameter settings (cell or nucleus intensity threshold, cell tracking duration) by Imaris will lead to different segmented cell images in a frame. This results in cell image data missing in some frames, thus our method allows a certain number of missing cell images in the selected uniform time periods and tries to find the maximum number of continuous cells images in this uniform time period.

Overall three cell image sets are generated for three phases, each with an approximately 1:5 ratio between progenitor cell images and normal MEFs cell images. For phase 1, 78 IPS progenitor cells and 391 normal MEFs are labeled; for phase 2, 84 IPS progenitor cells and 420 normal MEFs are labeled; for phase 3, 74 IPS progenitor cells and 370 normal MEFs are labeled. Each of these three initial cell image sets are divided into the training and test sets: 70% of cell images for each time phases are selected randomly as training set with the remainder (30%) as test set. The ratio between progenitor cell images and normal MEFs cell images is kept approximately 1:5 for these training and testing sets. For the the training sets, there are 55 iPS progenitor cells and 274 normal MEF cells in phase 1, 59 iPS progenitor cells and 294 normal MEF cells in phase 2, as well as 52 iPS progenitor cells and 259 normal MEF cells in phase 3.

In this paper, the initial cell dataset is used for cross-validating the proposed method, and the training dataset is used for missing value processing and feature selection. For different analytic steps, the specific data sample size depends on the time period from which the data has been collected. Numerical features are calculated for all cell images in the datasets and saved in CSV files. All datasets are standardized utilizing z-score.

### Missing values processing

Processing missing values for the cells in the corresponding frames is an important step for our model. Imaris cannot continuously identify all the cells in the frame due to different parameter settings or complex three-dimensional cell environment. This implies that there exists a certain number of cell images with missing feature values in the uniform time periods. A certain number of missing images in the frames are permitted for cells to guarantee a modest data size, and missing cell features are estimated with an imputation method. To choose the most appropriate approach, we first analyze the impact of the number of missing frames on the model, and then analyze the effect of three different imputation methods under the corresponding missing frame numbers. Details for the three imputation methods are as follows:

- *set_mean*. The missing value is set to the average value of all nonempty frames for a specific type of feature in its sample from the selected time window.
- *set_KNN*. The missing value is set to the weighted average value of five nearest nonempty neighbor frames for a specific type of feature in its sample. The calculation of weight uses k-Nearest Neighbor (KNN) algorithm. The formula is as

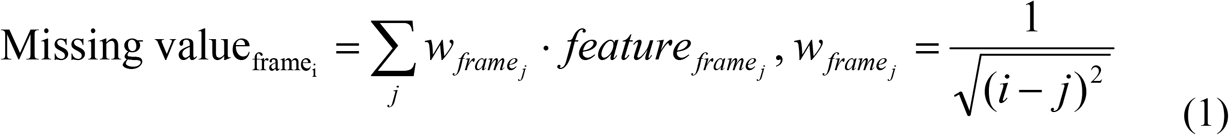

where *j* represents the index of five nearest frames neighbor for missing frame *i*.
- *set_mean_mod*. Missing value is set to the average value of five nearest nonempty neighbor frames for a specific type of feature in its sample.

### Time window and feature selection

Because of the two rules used in dataset generation (Section **Cell image datasets generation**), although images are provided up to 49 hours, it is unable to construct the model based on the whole period. From a total of 49 hours, numerous time periods can be chosen, and the model needs to select best time windows among all these eligible time periods. Time window selection includes start frame selection and window length selection. Start frame represents the moment that the time window starts from, and window length represents frame number that the time window contains. For each time window with a selected time frame and window length, we train and validate the proposed method on the corresponding dataset generated. Validation is performed with 5-fold cross validation and the evaluation metric is precision.

Morphological and motion feature selection is used to improve the performance. Since it is difficult to guarantee image recording time to be accurately consistent for every batch through experiments, model performance needs to be robust among wider time periods. Every type of features contains multiple frames of features from the corresponding best time windows. Features in a time window are treated as a bundle so we can learn the dynamic cell growth process.

There are two steps for feature selection. The first step is recursive feature elimination. Firstly, we use all 11 types of features to train the model with 5-fold cross validation and calculate its precision as initial unimportance score. Then we delete each type of feature at a time and obtain 11 precision values as new unimportance scores. We compare every new score with the initial score, and remove the feature type with the largest unimportance score higher than initial score. The recursive process will be repeated on feature set until the model performance can be no longer improved or there is no feature. We then rank the importance of all 11 types of features and delete the least important feature types. Second, we calculate the Pearson correlation coefficient for the selected feature types from step 1 to remove the highly correlated features with a correlation coefficient of 0.60 or above.

### Machine learning model and validation

XGBoost, a Boosting algorithm, is used in this paper for feature selection and IPS cell recognition. XGBoost integrates many weak tree-classifiers together to form a strong classifier. This algorithm applies numerous strategies to prevent overfitting, and it is widely utilized in data science such as cell analysis [39–43]. Hyperparameters of XGBoost are tuned using grid-search for model training with selected features and best time windows.

For model validation, firstly we use 5-fold cross-validation on the initial cell image datasets from the time windows of the three phases. Dataset generated from initial cell-sets contains about 70 iPS cells for each phase. The ratio of iPS cells and normal MEFs keeps as 1:5 in each dataset.

In order to test the model’s ability/robustness to predict the iPS progenitor cells around the neighborhood of the corresponding training time window, holdout validation is performed. Because iCD1 medium change is operated manually during the experiments, it is impracticable to guarantee that for per batch data the duration of medium change is accurately consistent with the existing data. This inconsistency might lead to a non-exact match between the timeline after medium change and the timeline used in the model training process. The holdout validation is designed as follows, for the model trained on time window *i~j*, we examine the model’s performance on several neighbor time windows, including time windows *i-3 ~ j-3, i-2 ~ j-2, i-1 ~ j-1, i ~ j, i*+*1 ~ j*+*1, i*+*2 ~ j*+*2*, and *i*+*3* ~ *j*+*3*, where *i* represents start time frame of the window and *j* represents the terminal frame. The training dataset from time window *i~j* is generated from the initial training image data sets (70% of the initial total dataset), and test datasets of the seven neighbor time windows are generated from the test datasets (30% of the initial total dataset).

Moreover, in order to further test our model’s ability to predict the iPS progenitor cell on a time window which doesn’t overlap with the window in the training process, an independent test is performed. Model performance is tested on time windows which are far away from the training time windows. Since we have three time phases, we first select test time windows in phase 2 and 3 for the models trained on time windows of phase 1 and 2 respectively. For testing our model developed for phase 3, we select the independent test time windows also in phase 3, but without any overlap with the corresponding training time windows.

### Evaluation metrics

In this paper, precision is mainly used for evaluation defined as,

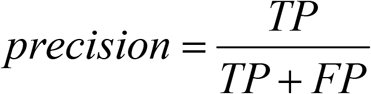

where TP and FP represent the number of true positive and false positive prediction. This metric evaluates the accuracy for the positive sample predicted by the model. Biologists need a cell sample set enriched with true iPS progenitor cells so that in the early stage of reprogramming progenitor cells can be studied with high probability.

## Results and Discussion

### Missing frames processing and imputation method

First, the effect of missing frames and imputation methods on the model’s performance was analyzed. Experiment for missing value was performed under six kinds of missing frame numbers, which were numbers below or equal to five, four, three, two, one and zero. Model performance was tested for each missing frame number with three imputation methods on time periods of two window lengths (10 and 19 frames) located in three phases, which were time period/window 19h30min ~ 21h10min from phase 1 (TP1), 25h50min ~ 27h30min from phase 1 (TP2), 41h50min ~ 43h30min from phase 2 (TP3), 18h10min ~ 21h20min from phase 2 (TP4), 26h ~ 29h10min from phase 3 (TP5) and 42h ~ 45h10min from phase 3 (TP6).

Two window lengths (10 and 19 frames) were selected because a reasonable number of continuous cell images could be traced. A short window will have more data but the motion and morphological pattern of iPS progenitor cells cannot be learned while a long window will result in a much smaller dataset. For each length, we chose three time windows randomly to study whether different lengths would affect model performance under uniform missing frame number. Datasets were generated from the training datasets, which were about 52~59 iPS cells and 259~294 normal MEFs for time windows with 10 frames, 43~50 iPS cells and 238~264 normal MEFs for time windows with 19 frames. Model was evaluated by the average precision with 5-fold cross validation over 20 times.

**Fig 2** showed the comparison results of different missing frame numbers and imputation methods. For each missing number and imputation method, **Fig 2(a)** described the average precision over six time windows (TP1 to TP6), indicated by blue boxes for set_KNN, red boxes for set_mean and green boxes for set_mean_mod. Also shown in **Fig 2(a)** was the average precision over all three imputation methods, indicated by grey boxes. **Fig 2(b)** described the standard deviations of the corresponding precision values in **Fig 2(a)**. Detailed precisions for all six time periods (TP1~TP6) were provided in Figure S1 of the **S1 File**.

**Fig 2.**
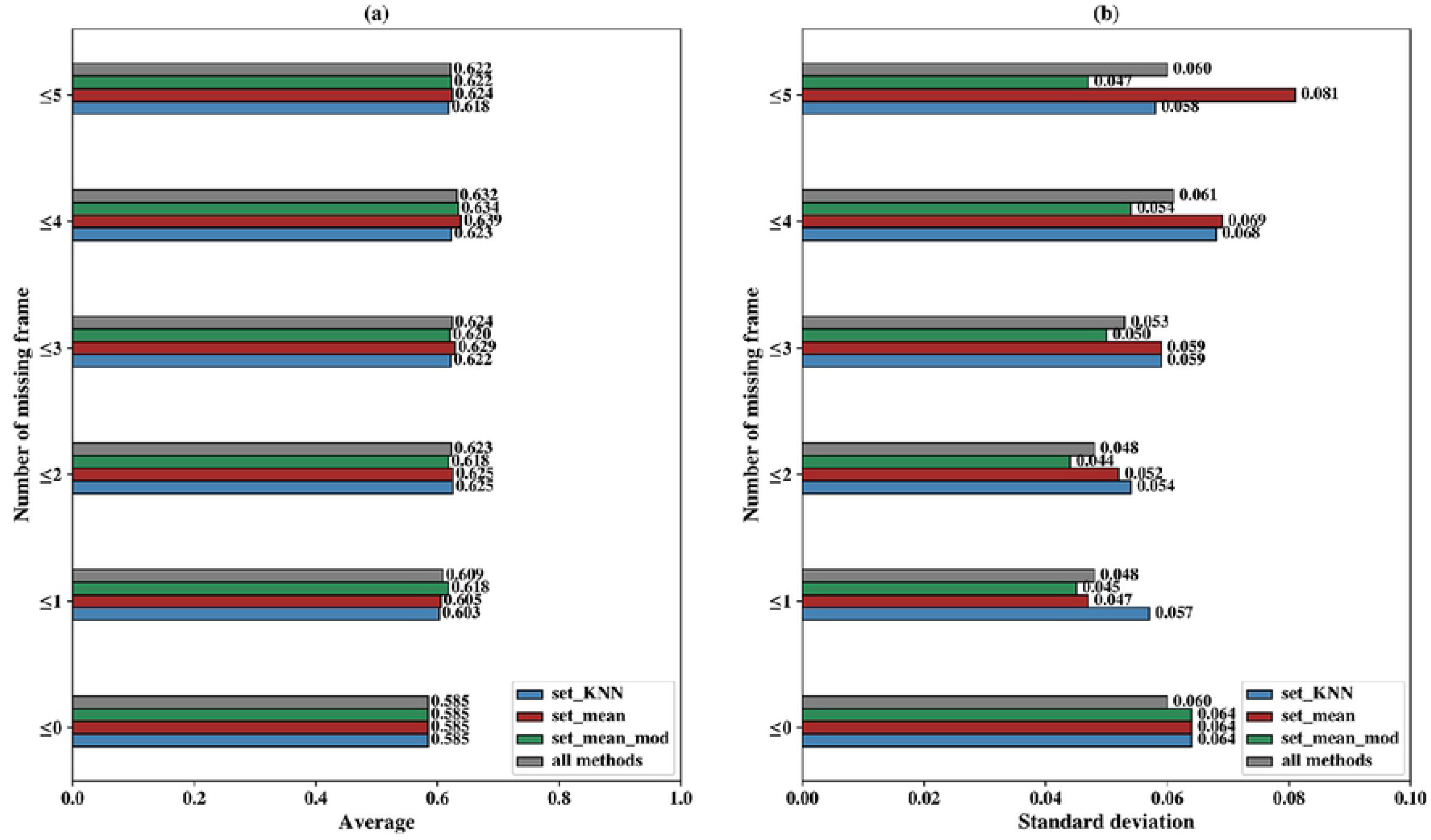
Model comparison for different missing frame number and imputation methods. Fig 2(a) shows the average precision over six time periods (TP1 to TP6) for each missing frame number and imputation method set_KNN (colored as blue), set_mean (colored as red), set_mean_mod (colored as green) and all three imputation methods (colored as gray). Fig 2(b) shows the standard deviation, as a function of missing frame number, of imputation method set_KNN (colored as blue), set_mean (colored as red), set_mean_mod (colored as green) and all three imputation methods (colored as gray).

**Fig 2(a)** showed that precision was higher when several missing frames were allowed. For missing frame number of 0, the average precision of all method was only 0.585 and all the average precisions of non-zero missing frame numbers were higher than 0.585. **Fig 2(a)** also showed that the maximum average precision of all method was about 0.632 under missing frames of 4, 4.7% higher than precision under no missing frames and 0.9% higher than precision under missing frame number of 2. On one hand, the size of the dataset is larger when missing value is permitted, on the other hand, the missing frame may introduce new pattern for classification because iPS progenitor cells proliferate more frequently than normal MEFs, and cell division can partly result in missing value. When cells divide at a certain frame in their time periods, the feature values of all subsequent frames are missing.

In **Fig 2(b)**, the maximum standard deviation of all methods as indicated by gray box was 0.061 under missing 4 frames. For each specific method, the maximum standard deviation was 0.081 for Set_mean under 5 missing frames. The precision with two missing frame numbers had the minimum standard deviation for all method (0.048 as indicated by gray boxes) and at the same time it was also very close to the maximum precision (0.623 compared with the maximum value of 0.632 in **Fig 2(a)**). In addition, Set_mean_mod showed the minimum standard deviation of all 3 imputation methods for all missing frame numbers (indicated by green boxes), an indication of stable performance. Although Set_mean_mod also showed smallest standard deviation for missing frame number of 1, its precision value of missing frame number was smaller than that of missing frame number of 2. Therefore, we used missing frame number less than or equal to two and select imputation method as set_mean_mod in our model.

### Time window selection

Time window selection was performed to select best time windows with high precision for each phase. Since Imaris could not detect all cell images in every frame, the whole time periods of three phases were divided into numerous time windows. For time window selection (including start frame and window length), we set start frame to 21 time points which were 18h20min, 18h40min, 19h, 19h20min, 19h40min, 20h, 20h20min, 26h10min, 26h30min, 26h50min, 27h10min, 27h30min, 27h50min, 28h10min, 42h10min, 42h30min, 42h50min, 43h10min, 43h30min, 43h50min, 44h10min in three phases. Meanwhile, we set window length to 12 different values including 7, 9, 11, 13, 15, 17, 19, 21, 23, 25, 27 and 29 frames. For the total of 252 (12 times 21) time windows, we first generated datasets for each time window with 11 types of morphological/motion features. All datasets were generated based on the training dataset and contained about 38~59 iPS progenitor cells and about 190~295 normal MEFs. Then we selected the optimal time window through 5-fold cross-validation based on 20 XGBoost runs.

The model performance on these different time windows was shown in **Fig 3**. In this figure, shorter window lengths were marked in red colors and longer window lengths were marked in blue colors. We observed that precision of longer window lengths was lower than that of shorter window lengths in three phases, and this trend was less pronounced for phase 3. The size of the dataset may be the major reason for this trend. Due to the two rules in dataset generation, the amount of samples satisfying conditions decreases gradually with the increasing window length. For window length of 29 frames, there are just about 38 iPS progenitor cells and 262 normal MEFs in phase 1, about 44 iPS progenitor cells and 283 normal MEFs in phase 2, about 36 iPS progenitor cells and 242 normal MEFs in phase 3. As compared with the window length of 7 frames, there are about 53 iPS progenitor cells and 290 normal MEFs in phase 1, about 55 iPS progenitor cells and 285 normal MEFs in phase 2, about 57 iPS progenitor cells and 300 normal MEFs in phase 3. On the other hand, the number of samples is much less for later start frame than that for previous time since some cells have divided. For instance, there are only about 30 iPS progenitor cells and 200 normal MEFs for the last start frame with length of 29 frames in phase 3.

**Fig 3.**
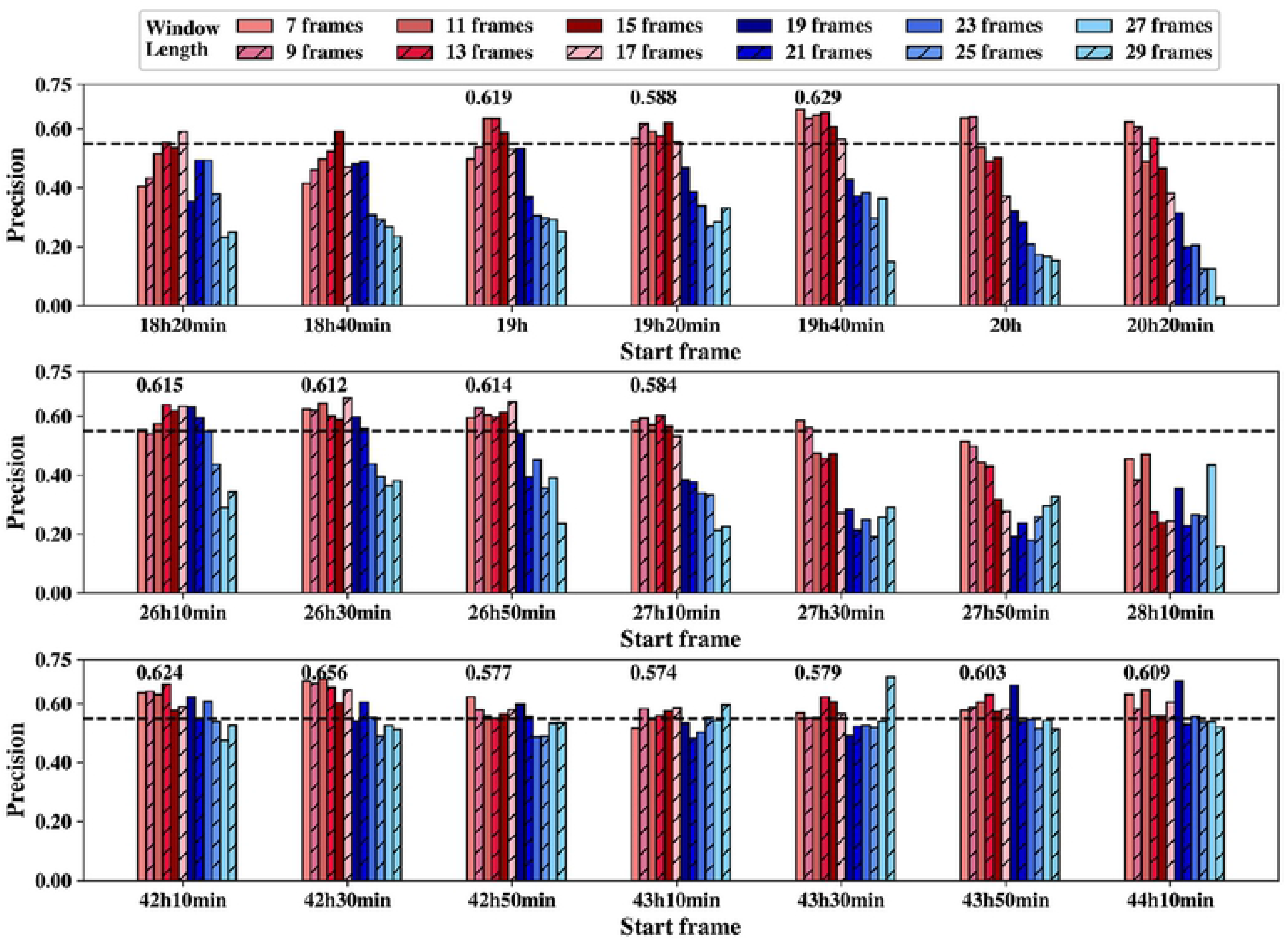
Time window selection. The three subplots represent the precision values for different time windows based on 21 start frames (x axis) and 12 window lengths (7 frames to 29 frames) for phases 1, 2, and 3 (from top to bottom) respectively, and the black bash line in each subplot indicates a precision value of 0.55.

Selection of best time windows according to maximum precision resulted in an unstable prediction performance. For instance, precision achieved the maximum value on the time window starting at 43h30min with length of 29 frames while all its adjacent time windows had poor performance with lower precision. It is unlikely to achieve the same performance on a new dataset of the same time window.

We selected the best start frame for each phase respectively. To exclude the start frame with high prediction precision for only 1 or 2 window lengths, 14 candidates of best start frames were selected when precision was above 0.55 for at least three successive window lengths. For each candidate best start frame, the average precision was calculated over the successive window lengths whose precision was above 0.55 and the average precision values were shown above each candidate best start frame in **Fig 3**. We only selected one best start frame for each phase according to the average precision values of the candidate best start frames, resulted in 19h40min, 26h10min and 42h30min for phases 1, 2 and 3, respectively.

Secondly, the candidate best window lengths were selected whose precision values were all above 0.55 for 3 best start frames of step 1, resulting in window lengths 11, 13, 15 and 17 frames. For each window length, the precision values, average precisions and the corresponding standard deviation of 3 different best start frames were provided in Table S1 of **S1 File**. The average precision of 0.640 for window length of 13 frame was the highest while its standard deviation was the smallest (0.01), thus window length of 13 frames was selected as the best window length.

### Two-step feature selection

We performed a two-step feature selection method on three phases respectively. Firstly, we generated datasets from best time windows based on the training cell image datasets. The dataset of each phase contained 11 types of morphological and motion features, all of which contained about 50~59 iPS progenitor cells and about 200~295 normal MEFs.

For the first step, an iterative feature removal procedure was performed on the corresponding dataset of each phase to study the importance of each feature type. Average precision was calculated via 5-fold cross-validation over 20 runs on the dataset of each phase, and later sets as initial unimportance score. Next, we removed each type of features and calculated the unimportance scores (average precision). Feature with maximum score would be deleted only if this score was greater than the initial unimportance score, which would then be updated as the maximum score. This step was repeated until no score was greater than initial score or no more feature could be selected.

Results from step 1 feature selection were shown in **Fig 4**. For phase 1 precision was no longer improving after removing ellipsoid-oblate, displacement and volume; for phase 2 precision was no longer improving after removing displacement and volume; for phase 3 precision was no longer improving after removing displacement, ellipsoid-prolate, area and volume. In the end, eight types of features were selected for phase 1, nine types of features were retained for phase 2, and seven types of features were retained for phase 3. Selected features from this step were indicated in **Fig 4** by star symbols. The corresponding precisions for best windows with 13 frames before feature selection were 0.624, 0.607, 0.646 for phases 1, 2 and 3, respectively, and after feature selection, these precision values had increased to 0.691, 0.613 and 0.682 respectively.

**Fig 4.**
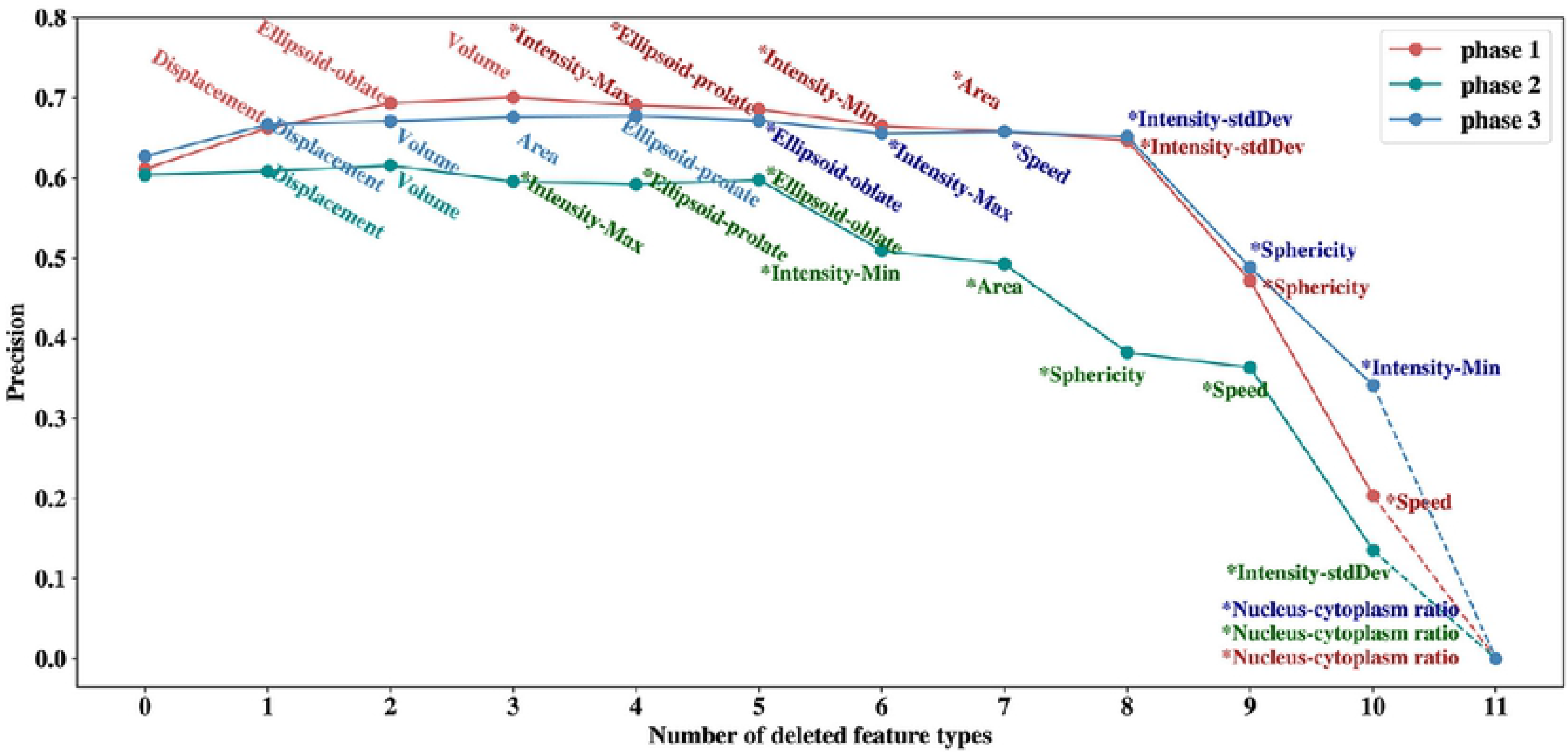
Feature ranking and selection. This figure shows how the precision values change with the deleted feature in a recursive fashion. Least important features are removed earlier.

The removing order of feature type in **Fig 4** indicated the importance of each feature type. We observed from **Fig 4** that three types of features, nucleus-cytoplasm ratio, sphericity and intensity-StdDev, were important among all three phases. Nucleus-cytoplasm ratio was the top important factor in three phases. Sphericity and intensity-StdDev were among the top 4 common features of three phases. Intensity showed clear different patterns between normal MEFs and progenitor cells. As shown in **Fig 5(a)**, the progenitor cells in the blue circles showed a uniform intensity distribution between nucleus and cytoplasm, while for normal MEFs in the yellow boxes, the cytoplasm showed weaker intensity as indicated by the blurring edges. Also shown in **Fig 5(a)**, the nucleus and cytoplasm of progenitor cells in the blue circles and normal MEFs in the yellow boxes were enlarged and colored by light blue and green respectively. It is clear that nucleus-cytoplasm ratio for progenitor cells are much larger than that of normal MEFs. From **Fig 5(a)**, the cell area of progenitor cells is also smaller on average than normal MEFs, indicating the importance of sphericity since area is closely related to sphericity by the equation from Part 1 of the S1 File. The selected features are consistent with the experimental results that iPS progenitor cells exhibit higher nucleus-cytoplasm ratio, smaller total area, and higher proliferation rate than normal MEFs[21].

**Fig 5.**
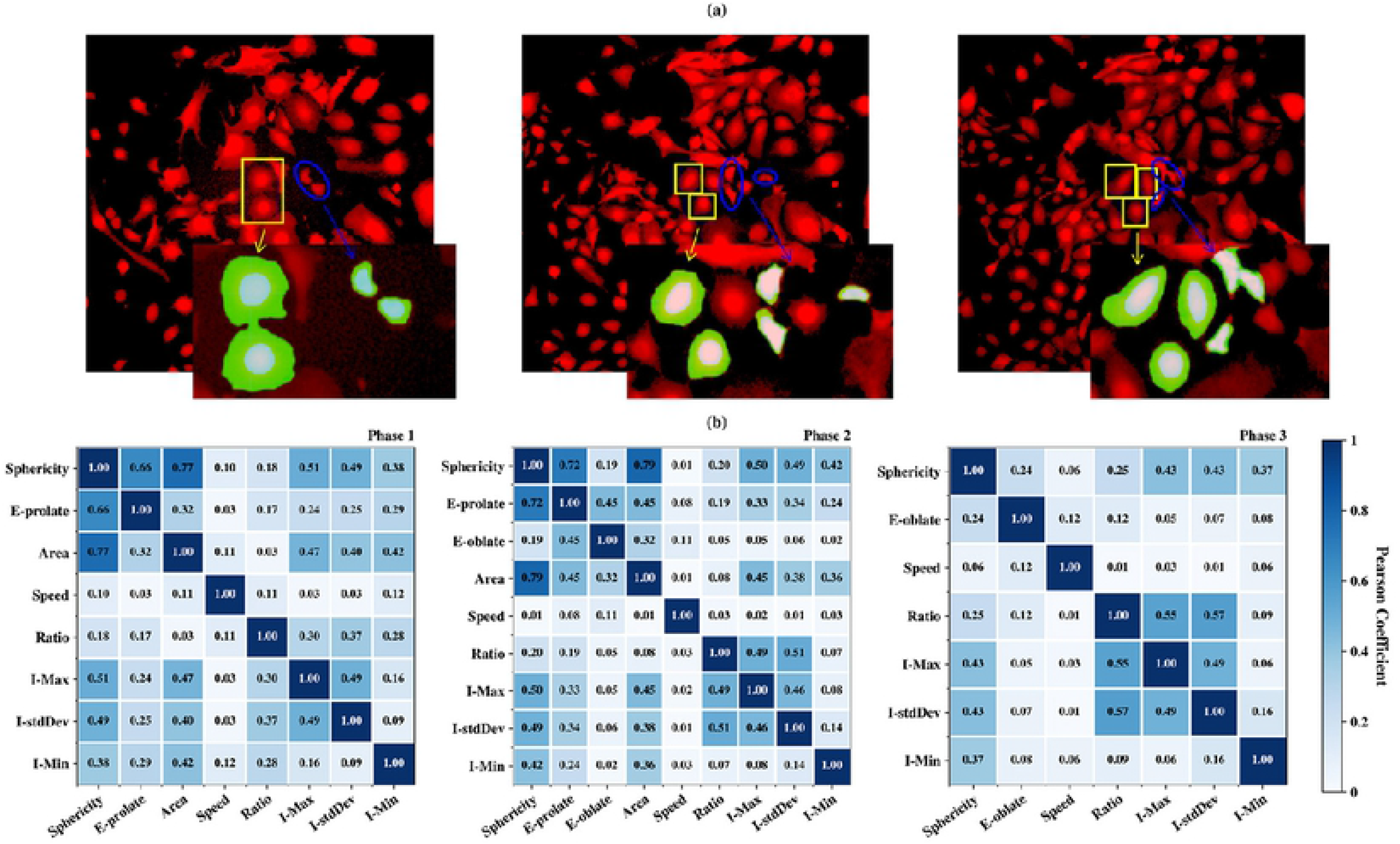
iPS progenitor cells vs. MEFs and Feature correlation. (a) shows the examples of iPS progenitor cell images (blue circles) and normal MEFs images (yellow boxes) taken from phase 1, 2 and 3 of field 2 (Left, middle and right). Nucleus and cytoplasm of the enlarged progenitor cells and normal MEFs are colored in light blue and green respectively. (b) shows the Pearson coefficients between remaining types of features in three phases after the first step of feature selection. Note in this figure ellipsoid-prolate is denoted as E-prolate, intensity-StdDev as I-stdDev, intensity-min as I-Min, intensity-max as I-Max, nucleus-cytoplasm volume ratio as Ratio, ellipsoid-oblate as E-oblate.

In order to further study the correlations of different features, as a second step we calculated the Pearson correlation coefficients between the selected features. The results for three phases were shown in **Fig 5(b)**. In our model, two feature types were considered strongly correlated if the coefficient was greater than 0.6 and one of them was removed. When two different feature types were strongly correlated with a third feature type, both of them were removed with the purpose of keeping as less number of features as possible. For phase 1, the coefficient between sphericity and area was 0.77 in phase 1, and the coefficient between sphericity and ellipsoid-prolate was 0.66, thus area and ellipsoid-prolate were removed from the list. Similarly, they were removed for phase 2 as well. The strong correlation between sphericity, ellipsoid-prolate and area is caused by the fact that Imaris extracts features from two-dimensional cell images assuming cell thickness as constant. Furthermore, since ellipsoid-oblate was associated with cell thickness, it was removed from the feature list as well for phase 2 and phase 3. Overall, six types of features (Sphericity, I-Min, I-stdDev, I-Max, Ratio, Speed) were selected for all the models.

### Cross-validation

With selected features, a grid-search scheme was used for hyperparameter optimization of XGBoost with 5-fold cross-validation, and the datasets were generated based on the training sets for three phases. Three hyperparameters such as learning_rate, n_estimators and gamma were set to 0.01, 385 and 0 respectively. We had validated our model with three different experiments as shown in **Fig 1**.

For cross-validation, datasets were generated from initial whole cell image dataset. Dataset for phase 1 contained about 63 iPS progenitor cells and about 326 normal MEFs. Dataset for phase 2 contained about 82 iPS progenitor cells and about 427 normal MEFs. Dataset for phase 3 contained about 72 iPS progenitor cells and about 359 normal MEFs. For each phase, 5-fold cross validation was performed 10 times on every best time windows with 6 selected feature types, resulting in a total of 117 for window length of 13 frames. **Fig 6(a)** showed precision scores for 3 different phases, and all of the precision values were above 0.580. For phase 1, the precision value was highest, 0.732.

**Fig 6.**
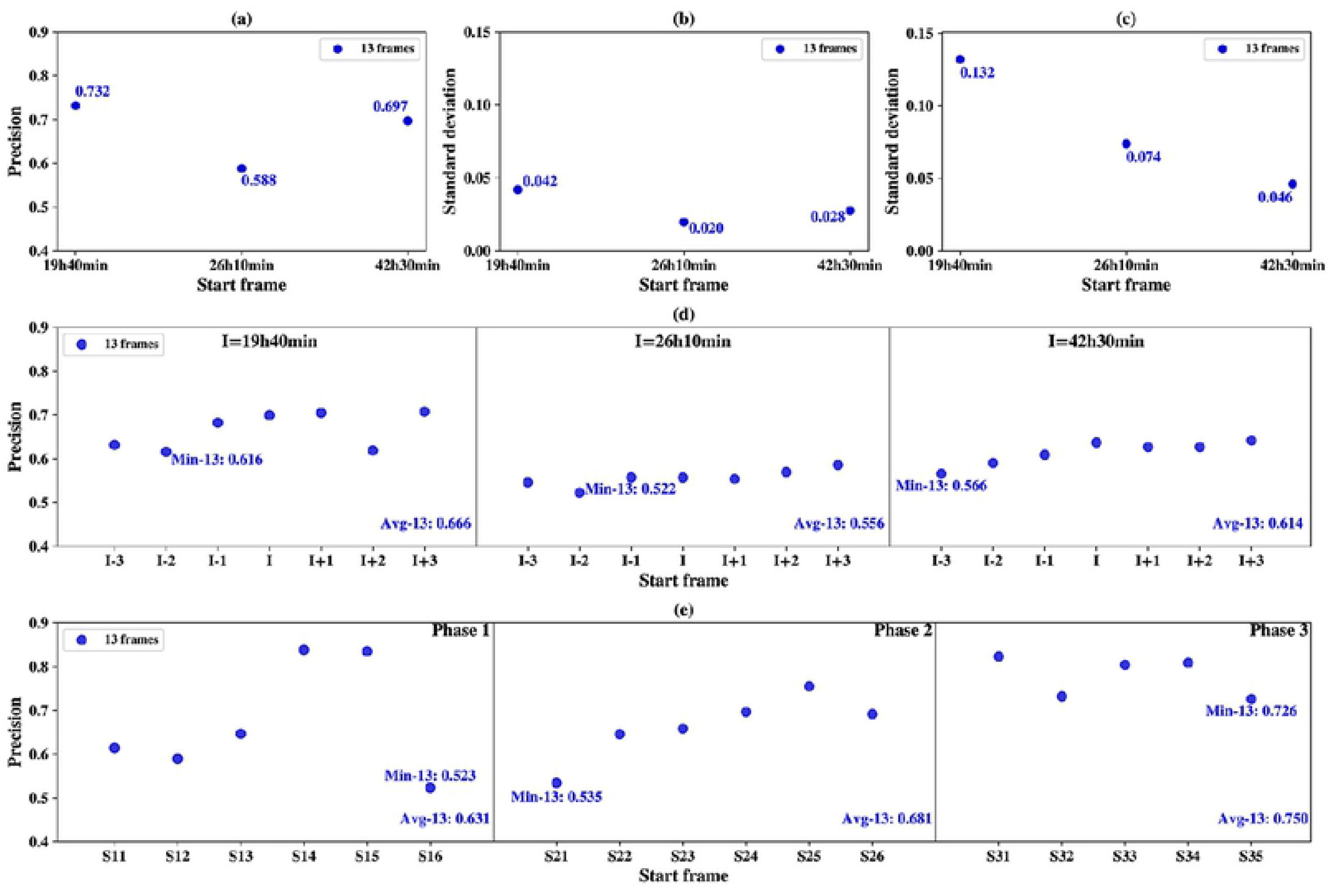
Model validation. In all sub-figures, X axis indicates the start frame of the best time windows and the corresponding window length (13 frames) is indicated in the inlet. (a) 5-fold cross-validation precisions over 10 runs. (b) the standard deviation of the average precision of the neighborhood time windows in Figure 6(d). (c) the standard deviation of the average precision of the distant windows in Figure 6(e). (d) the average precision of seven neighborhood time windows calculated over 10 holdout validation runs. (e) the average precision over 10 independent tests for six best time windows on their corresponding distant windows.

### Holdout validation

Holdout validation was used to test the model’s ability to predict the iPS progenitor cells in the neighborhood of the time window in which the model had been trained. Since in real application, it is difficult to generate the dataset whose images have the exact start time as in the training dataset, holdout-validation is very important for testing the model’s generality on the neighborhood time windows. For each phase, the training dataset for window length of 13 frames was generated. In phase 1, the window start frame I was 19h40min as shown in **Fig 6(d)**. Models trained on this dataset was then tested on seven test datasets corresponding to start frames I, I-1, I-2, I-3, I+1, I+2 and I+3, illustrated in **Fig 1** and **Fig 6(d)**. There was no overlap between the training and testing datasets.

For each time window, average precision value was computed over 10 holdout validation runs, and the results were shown in **Fig 6(d)**. The minimum average precision values were 0.616 for window length of 13 frames and start frame I-2 in phase 1, 0.522 for window length of 13 frames and start frame I-2 in phase 2 and 0.566 for window length of 13 frames and start frame I-3 in phase 3. These minimum precisions were all smaller than the corresponding precisions in **Fig 6(a)**; what is more, **Fig 6(d)** also showed the average precision values for phase 1, 2 and 3 were all smaller than the cross-validation resulted in **Fig 6(a)**, indicating the difficulties for predicting the neighborhood time windows.

For each result of the 3 phases in **Fig 6(d)**, the standard deviations of average precisions were computed for window length of 13 frames in **Fig 6(b)**. The maximum deviation was 0.042 for window length of 13 frames in phase 1 and this indicated the trained models were relatively stable in terms of prediction precision in a wide range of neighborhood windows.

### Independent test

Finally, to test the model’s ability to predict the iPS progenitor cells on a distant time window without overlapped frames with the training window, we performed an independent test. If the training cell trajectory is long and contains enough typical iPS progenitor cells, the trained model on one window should be able to identify the motion and morphological patterns of iPS progenitor cells against normal MEFs, regardless of the selected time window.

For phase 1, the model trained on time window 19h40min~21h40min (length of 13 frames) was tested on time windows of phase 2, including time windows starting from 26h20min (S11), 26h40min (S12), 27h (S13), 27h20min (S14), 27h40min (S15), and 28h (S16), shown in the first panel of **Fig 6(e)**. Similarly, for phase 2, the model trained on time windows 26h10min~28h10min (length of 13 frames) was tested on six time windows of phase 3 starting from 42h10min (S21), 42h30min (S22), 42h50min (S23), 43h10min (S24), 43h30min (S25), 43h50min (S26), shown in the middle panel of **Fig 6(e)**. Lastly, for phase 3, model testing was performed on the distant time windows without overlapped frames from the same phase, shown in the right panel of **Fig 6(e)**. For time windows 42h30min~44h30min, we selected test time windows starting from 45h10min (S31), 45h30min (S32), 45h50min (S33), 46h10min (S34), 46h30min (S35).

Results of the independent test runs were shown in **Fig 6(e)**. The minimum precision was 0.523 for window length of 13 frames for S16 in phase 1. The average precision of phase 1 was lower than those of holdout validation and cross-validation, however, the average precision of phase 2 and 3 were both better than cross-validation and holdout validation. For the prediction of distant time windows, our model could have worse performance than that of neighborhood windows, but our model could also outperform the cross validation and holdout validation (indicated by the standard deviation in **Fig 6(c)**). The reason was the independent test datasets for phase 2 and 3 were closely related to the training dataset. The standard deviations of the independent tests were much higher than those of the holdout validation, which could also be seen from the large fluctuations of the precision values in **Fig 6(e)**. Nevertheless, the minimum average prediction precision was above 52% among all the experiments, and maximum average precision was about 0.750 for the independent test in phase 3.

## Conclusion

In this paper, we proposed a machine learning based model together with time-lapse image analysis to predict/identify iPS progenitor cells during the first 3-5 days after reprogramming initiation. The model generated a variety of morphological and motion features among different time windows, then relied on a two-step feature selection algorithm to select the most important features. The proposed computational approach is very unique from previous experimental techniques which identify the iPS progenitor cells by retrospectively tracking the cell images manually frame by frame from the image frame of GFP expression.

By the experimental study of the enriched iPS progenitor cells in the early stage of reprogramming, the proposed method could provide a new technique or attempt for experimenters to improve the iPS reprogramming efficiency and to study the underlying mechanism of iPS reprogramming. Morphological and motion features, especially sphericity, intensity-StdDev and nucleus-cytoplasm volume ratio, have been found most important for the progenitor cell classification, which is consistent with the experimental observations.

Cross-validation of the proposed method trained and tested on the same time window showed that the prediction precision is above 0.580 for all three phases. Since in real applications, it is very difficult to match imaging timeline precisely between different experiments, holdout validation and an independent test are also performed to test the model’s ability to predict iPS progenitor cells in the neighborhood time windows and distant time windows, respectively. The results showed our model can predict the iPS progenitor cells with a minimum precision of 52% for neighborhood windows and distant windows, and the maximum average precision is about 0.750 for the independent test in phase 3. The prediction performance of our model tends to have a larger fluctuation for distant windows than for neighborhood windows, indicated by the larger standard deviation of independent test runs.

For future works, models on different time windows for each phase can be combined to achieve higher prediction accuracy.

## Acknowledgment

This work is supported by the National Key Research and Development Program of China under Grant No. 2016YFB0201305 and 2018YFB0204403; National Science Foundation of China under grant no. U1435215 and 61433012; the Shenzhen Basic Research Fund under grant no. JCYJ20160331190123578, JCYJ20170413093358429 and GGFW2017073114031767; Chinese Academy of Sciences grant under no. 2019VBA0009. We would also like to thank the funding support by the Shenzhen Discipline Construction Project for Urban Computing and Data Intelligence, Youth Innovation Promotion Association, CAS to Yanjie Wei.

## Supporting information

**S1 File. Supplementary Material**

